# Leveraging pathogen community distributions to understand outbreak and emergence potential

**DOI:** 10.1101/336065

**Authors:** Tad A. Dallas, Colin J. Carlson, Timothée Poisot

## Abstract

Understanding pathogen outbreak and emergence events has important implications to the management of infectious disease. Apart from preempting infectious disease events, there is considerable interest in determining why certain pathogens are consistently found in some regions, and why others spontaneously emerge or reemerge over time. Here, we use a trait-free approach which leverages information on the global community of human infectious diseases to estimate the potential for pathogen outbreak, emergence, and re-emergence events over time. Our approach uses pairwise dissimilarities among pathogen distributions between countries and country-level pathogen composition to quantify pathogen outbreak, emergence, and re-emergence potential as a function of time (e.g., number of years between training and prediction), pathogen type (e.g., virus), and transmission mode (e.g., vector-borne). We find that while outbreak and re-emergence potential are well captured by our simple model, prediction of emergence events remains elusive, and sudden global emergences like an influenza pandemic seem beyond the predictive capacity of the model. While our approach allows for dynamic predictability of outbreak and re-emergence events, data deficiencies and the stochastic nature of emergence events may preclude accurate prediction. Together, our results make a compelling case for incorporating a community ecological perspective into existing disease forecasting efforts.

## Introduction

The emergence of infectious diseases in humans and wildlife is a continuous and natural process that is nevertheless rapidly intensifying with global change (1). Around the world, the diversity, and frequency, of infectious outbreaks is rising over time (2; 1), and the vast majority of pathogens with zoonotic potential still have yet to emerge in human populations, with an estimated 600,000 minimum viruses with zoonotic potential (3). Intensifying pathways of contact between wildlife reservoirs and humans, and rapid spread of new pathogens among human populations around the globe, are considered major drivers in this accelerating process (4; 5). Changes in climate and land-use, as well as food insecurity and geopolitical conflict, are expected to exacerbate feedbacks between socio-ecological change and emerging infectious diseases (EIDs). In the face of these threats, the anticipation of disease emergence events is a seminal but elusive challenge for public health research (6).

One forecasting approach recognizes that the drivers of emergence events are distributed non-randomly in space and time, and follow predictable regional patterns that inherently predispose some areas to a higher burden of EIDs (7). Different classes of emerging pathogens (e.g., new pathogens versus drug-resistant strains of familiar ones; vector-borne and/or zoonotically transmitted diseases) follow different spatial risk patterns at a global scale (1). In part, this can be explained by the non-random distribution of host groups that disproportionately contribute to zoonotic emergence events, like bats and rodents (8; 9), and are likely to continue to do so (10; 11; 12). However, additional factors are strongly associated with the distribution of emerging infection risk; notably human population density, land cover, and land use change (7). In addition to these factors, deterministic emergence of disease is influenced by social, cultural, and economic factors (13; 14; 15; 16; 17).

As a consequence of this heterogeneity in host distributions and other contributing factors, emerging pathogens may follow Tobler’s First Law (“near things are more related than distant things”; (18)), and fall into a handful of global biogeographic regions with similar pathogen communities (19). However, with increasing global connectivity, both pathogens and the free-living organisms that host them are spreading around the world at an accelerating rate, and consequently the spatial structure of pathogen diversity is becoming less pronounced. One study examining a global pathogen-country network showed that modularity is decreasing while connectance is increasing over time: pathogen ranges are on average expanding, and over time, geographically-separate regions are facing more threats (20; 21). This process of biotic homogenization has critical implications for public health, as known diseases can become unfamiliar problems in novel locations, or can re-emerge in landscapes from which they were previously eradicated.

Leveraging disease ecology in global health settings requires models that consider disease emergence as a long-term process over space and time, extending beyond initial spillover events. Work that models the impact of human mobility networks has arisen out of the pandemic influenza literature (22; 23; 24), and has recently been successful in developing a multi-scale approach to anticipating emergence risk for hemorrhagic viruses in Africa (25). However, conceptually-similar work capable of modeling numerous pathogen species at large spatial scales is presently undeveloped. It has been suggested that countries who share pathogens might be more likely targets during a given pathogen outbreak (19), but this approach does not leverage information on the identity of the shared pathogens. Given the inherent need in estimating outbreak potential, and the current availability of data on outbreak events, there is a current pressing need to leverage existing data on numerous pathogen species to allow for dynamic prediction of potential pathogen outbreak or emergence events.

Here, we examine the predictability of pathogen biogeography over time using a similarity-based approach that utilizes data on all pathogen outbreaks in all countries, but does not require information on pathogen traits or spatial structure. In the process of modeling outbreak predictability, we test a basic but important hypothesis: do recurring outbreaks have a more predictable signal than emergence events (and, implicitly, are emergence events predictable)? Within emergences, we further note the subtle difference between emergence and re-emergence, and hypothesize the factors driving these might be subtly different. While both may be driven by genetic shifts in pathogens or changing land use patterns enhancing transmission risk, re-emergence events are more likely to be related to weakened healthcare infrastructure, prematurely-terminated eradication campaigns (26; 27), or low detection long-term persistence of environmental pathogen reservoirs (e.g., anthrax spores in the soil; (28)).

Finally, we examine whether pathogens show any differences in predictability based on agent, class, or transmission mode. Diseases of zoonotic origin (i.e. with animal hosts) and with vector-borne transmission might be harder to predict due to hidden constraints on their distribution and more complicated outbreak dynamics than directly-transmitted pathogens have. On the other hand, commonalities between species that share vectors or reservoir hosts might lead to similarities in distributions (a common notion in pathogen biogeography, as in how dengue models were frequently used in the early days of the Zika pandemic, given the shared vector *Aedes aegypti*; (29; 30)). In this case, community-based prediction could be more powerful for zoonotic and vector-borne diseases. Differential frequency of zoonotic and vector-borne transmission might also make different pathogen classes (viruses, bacteria, fungi, and macroparasites) more or less predictable, as might different dispersal ability on a global scale, with respiratory viruses usually presumed to spread the fastest, and macroparasites generally treated as the most dispersal-limited. Understanding how the role of community structure changes for these different pathogens can help contextualize the method we use, and understand how it might be built upon to account for these differences.

## Methods

### Pathogen emergence data

Data from the Global Infectious Diseases and Epidemiology Network (GIDEON) contains pathogen outbreak information at the country level obtained from case reports, governmental agencies, and published literature records (31; 32). Records with multiple etiological agents (e.g., *“Aeromonas* and marine *Vibrio* infx.”) and unresolved to agent level (e.g., “Respiratory viruses - miscellaneous”) were excluded from the model. In a handful of cases, we kept divisions between clinical presentations from the same pathogens, like cutaneous versus visceral leishmaniasis. The data obtained were yearly records between 1990 and 2016, and consisted of pathogen outbreak and emergence events for 234 pathogens across 224 countries. While there are some data for pathogen events between 1980 and 1990, the number of pathogen events reported was fewer than from 1990 onward, suggesting some potential reporting or sampling bias in these earlier years. Therefore, we restrict our analyses to pathogen occurrences after 1990. Based on supplemental data from (20) and updated with recent literature given several misclassifications, each was manually classified as a bacterial, viral, fungal, protozoan, or macroparasitic disease, and as vector-borne and/or zoonotic or neither. In some rare cases, these were left as unknown; for example, Oropouche virus is vector-borne but its sylvatic cycle remains uncertain, while the environmental origin of Bas-Congo virus is altogether unknown.

While much can be gained by leveraging data on multiple pathogens to predict outbreak or emergence potential, there are some drawbacks. The most pronounced is that pandemic events may strongly influence model predictions, such that a pandemic of one pathogen will decrease model performance when attempting to predict outbreak or emergence potential of other pathogens. We explore this further in the supplement, where we see the inclusion of influenza and the corresponding 2009 flu pandemic noticeably affects our model performance. As such, we remove influenza from the main text analyses, and place analyses containing flu in the supplement for comparison.

We distinguish between three different types of pathogen events; outbreak, reemergence, and emergence. Outbreaks are pathogen events are recurrent pathogen events, quantified as having occurred in a given country within three years of a given year. Re-emergence events are those that did not occur within three years, but have occurred at some time in a given country in the past (a cutoff we chose inspired by World Health Organization guidelines for certifying regional eradica-tion of poliovirus or dracunculiasis). Lastly, emergence events were considered as the first record of a pathogen within a country.

### Model structure

We developed a dissimilarity-based approach to forecast pathogen outbreak and emergence events that does not require country-level or pathogen traits data. Applying tools from community ecology, we calculated mean pairwise dissimilarity (Bray-Curtis index, 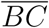) values for countries (how dissimilar are the pathogen communities between countries) and pathogens (how dissimilar are the geographic distributions of pathogens). For a given pair of countries *a, b* with *P*_*a*_ and *P*_*b*_ pathogens each, and *S* shared pathogens among those, the Bray-Curtis index is given as:

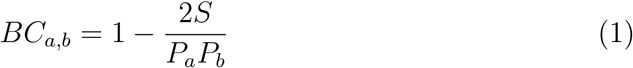

This can be treated as a measure of dissimilarity between different countries’ pathogen communities. We then considered the potential for a pathogen to be found in a country proportional to the product of these dissimilarity values. We also included year as a covariate, resulting in a set of four variables for model training.

Using these data, we applied a statistical approach previously used for species distribution modeling (33) and link prediction in ecological networks (34) called `plug-and-play` (PNP). This approach utilizes information on pathogen occurrence events, and also on background interactions — country-pathogen pairs which did not have a recorded outbreak — to estimate the suitability of a country for pathogen emergence from a particular pathogen (Figure 1). These suitability values can then be used to quantify model performance on data not used to train the model.

**Figure 1:**
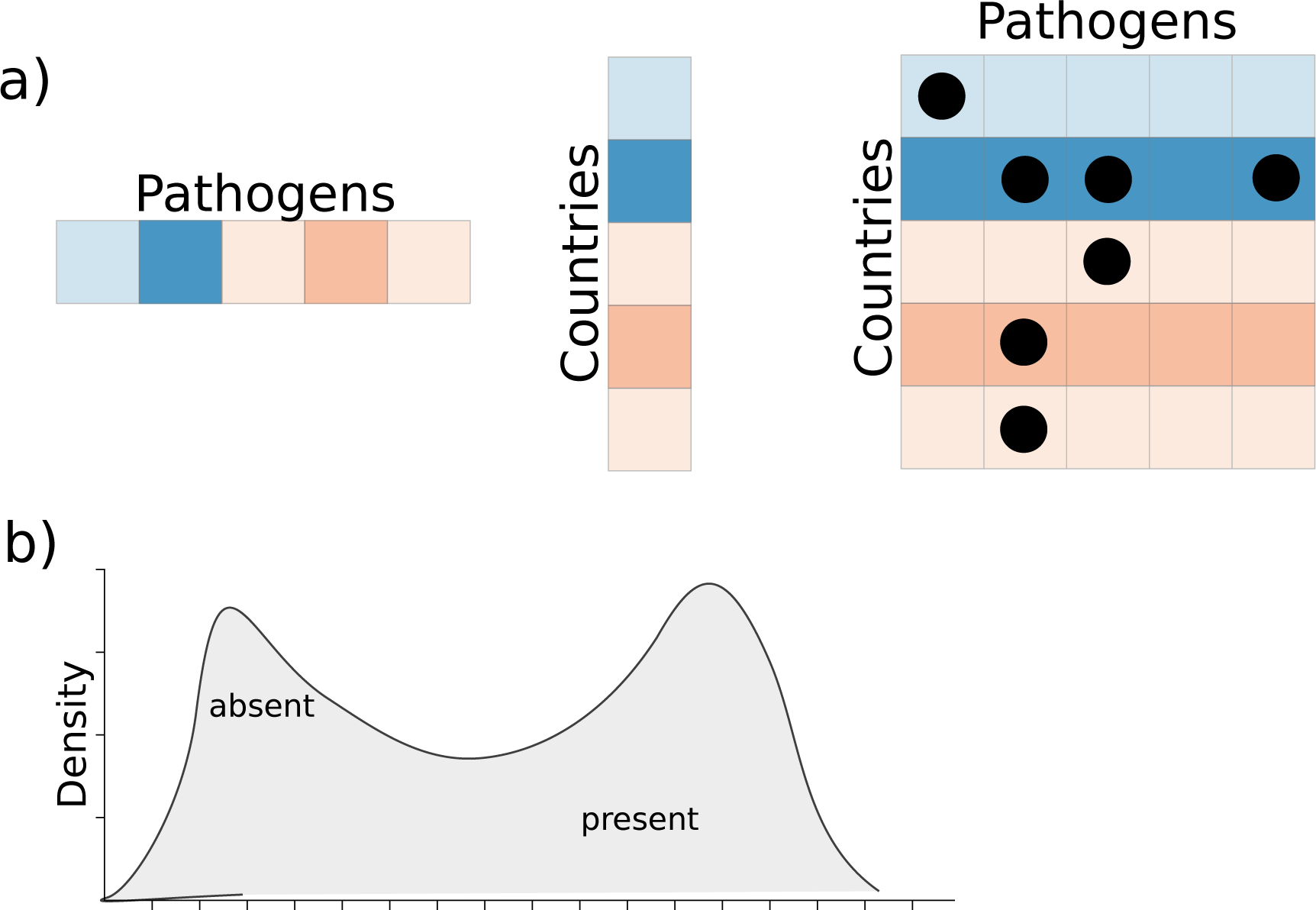
The dissimilarity-based model used takes mean dissimilarity values of pathogen distributions and between countries in a given year, and uses this information in addition to the product of these two values to train the PNP model. Pathogen occurrences among countries are present or absent (black dots in panel a indicate pathogen occurrences), and the density of dissimilarities where the pathogen occurred relative to the overall density of dissimilarities provides information on the suitability of pathogen occurrence in a given country (b), and forms the basis of the PNP model approach.

If pathogen outbreak events occur in the same countries probabilistically based on some propensity for the pathogen to occur at that location, we might expect that using past data on pathogen outbreaks could be used to forecast pathogen events. If a pathogen were to occur in a given country in one year out of four, a naive assumption would be that it has a 25% chance of occurrence in the subsequent sampling event. We examined how this null expectation compares against our approach, which uses information on country and pathogen similarity values (light grey lines in figures).

### Assessing model performance

We used the PNP modeling approach to address the possibility of predicting pathogen outbreak and emergence events compared to a null model. Model performance was quantified using Area Under the Curve (AUC), which captures the ability of the classifier to rank positive instances higher than negative instances. To assess model performance, we examined three different potential scenarios.

First, we examined how the inclusion of pathogen events from previous years influenced model accuracy. That is, we predicted pathogen events of 2016 using data starting at 2015 and then including additional years until 1995. This was performed to determine the amount of data necessary to make accurate forecasts. Second, we examined how predictive accuracy was maintained as we attempted to predict both past (hindcast) and future (forecast) pathogen events. To do this, we trained models on a ten year period (either 2005-2015 for hindcasting, or 1990-2000 for forecasting), and used these models to predict pathogen events between 1990 and 2004 for hindcasting, and between 2001 and 2015 for forecasting. Lastly, we examined how the accuracy of predictions might have changed over time. Given increased surveillance in more recent years, predictive accuracy might be dependent on the time period at which models are trained and predictions made. To test this, we trained models along a rolling window of 4 years from 1990-2015, using these models to predict pathogen events in the year following the final year of model training (e.g., a model trained on 1990-1994 would be used to predict pathogen events in 1995).

## Results

We find that our dissimilarity-based model can predict outbreak events accurately, re-emergence events slightly less accurately, and emergence events only slightly better than random (i.e., AUC = 0.5). This makes intuitive sense, as outbreak events occur repeatedly, providing not only ample data for model training, but also a clear tendency of a pathogen to occur in a country. That is, if the model is allowed to see 5 years of data, and the country has an outbreak of a particular pathogen in 4 of the 5 years, a naive model would predict that an outbreak will likely occur with an 80% probability. This situation corresponds to the null model (Figure 2), which performed poorly until enough temporal data was available, at which time the naive null model still underperformed our approach. Meanwhile, emergence events are determined by many unique drivers (7), which may not be consistent across any two given emergence events, and which we evidently lack sufficient data to predict using our method. While our model allows for dynamic predictability of outbreak and re-emergence events, data deficiencies and the stochastic nature of emergence events may thus preclude accurate prediction. However, our approach outperforms the naive null model in all modeled scenarios, especially when temporal data were limited (Figure 2–4).

**Figure 2:**
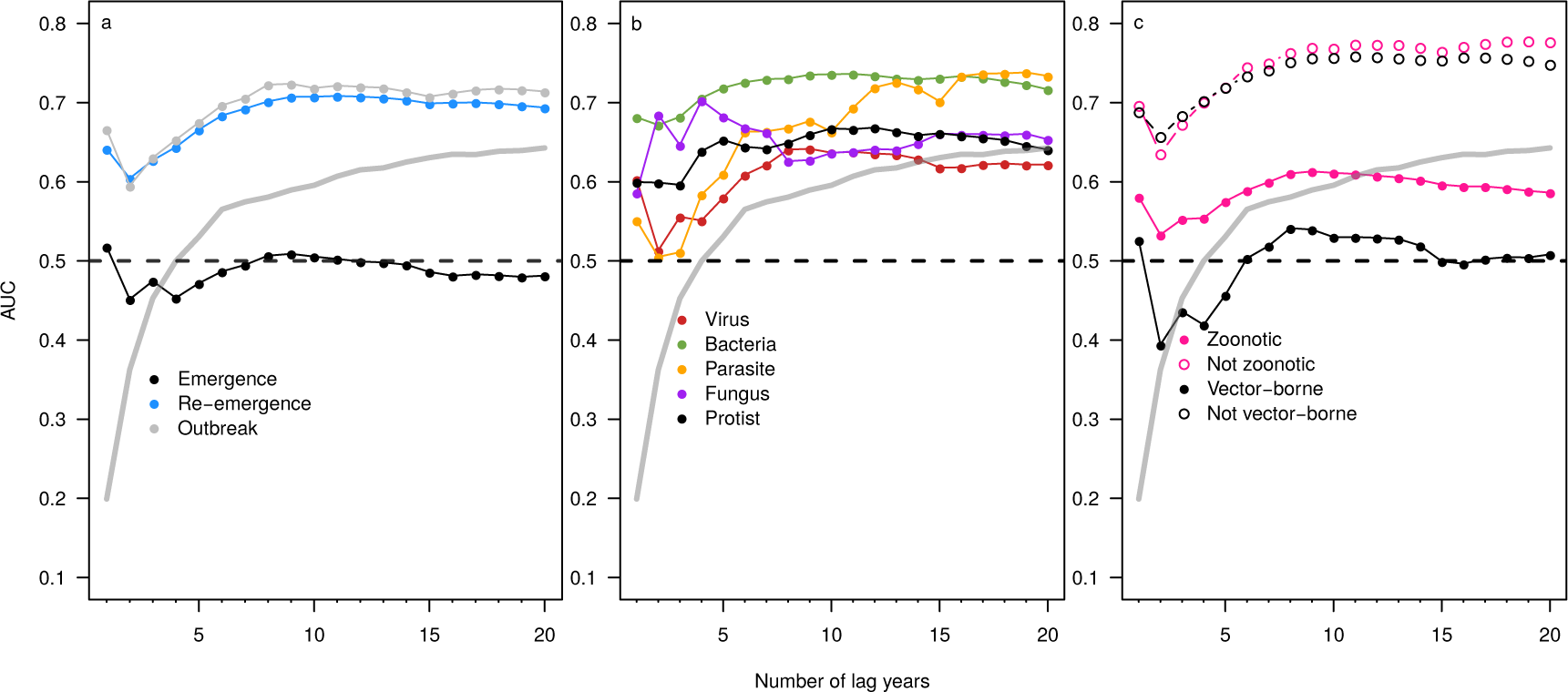
Pathogen events from previous years increased model predictive accuracy after an initial small decrease, suggesting that five years or more of data improves predictions, but accuracy could actually decrease in some data sparse situations where only two or three years of data were available. Performance of the null expectation (grey line) was less than our approach, except when the null was given more than 15 years of previous data.

**Figure 3:**
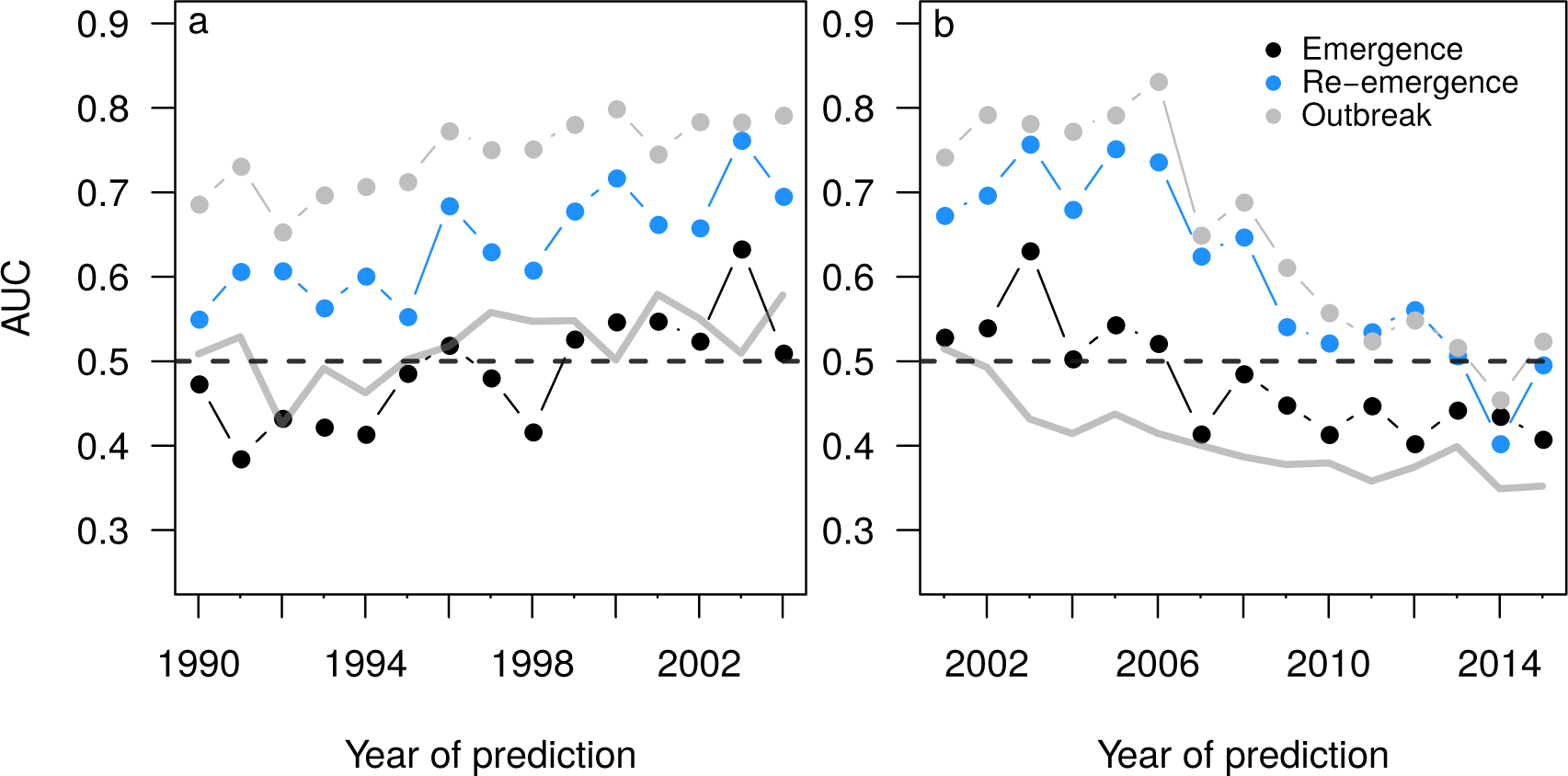
Predictive accuracy decreased when attempting to forecast far into the past or future. Models were trained on either the period between 2005-2015 (for prediction into the past) or 1990-2000 (for prediction into the future). The null expectation (grey line) performed consistently worse than our approach.

**Figure 4:**
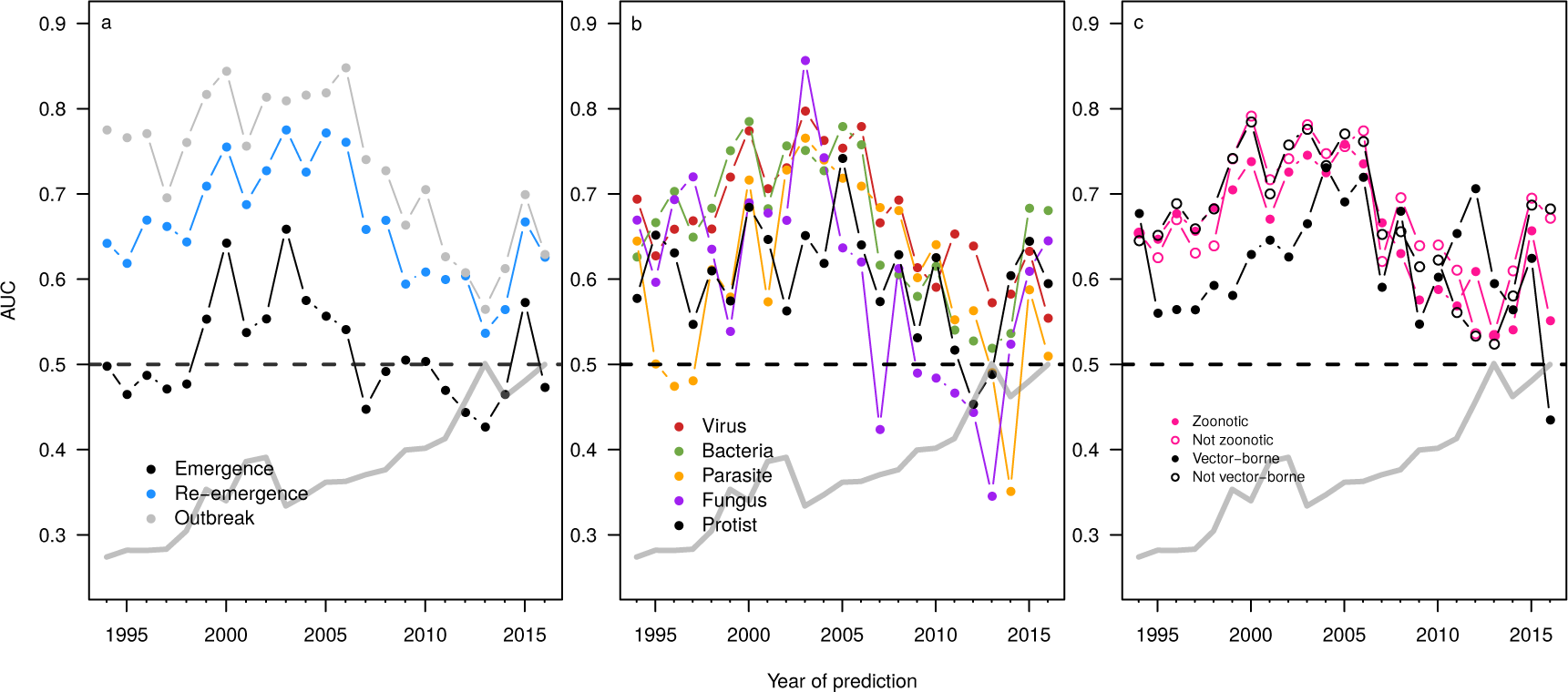
Using a rolling window (*t* = 4 years), we found that predictive accuracy did not increase as a result of enhanced surveillance and data collection of more recent years. The null expectation (grey line) performed consistently worse than our approach.

Our predictive model was sensitive to the number of training years (Figure 2), with accuracy plateauing around 5-10 years of training data; however, models also just trained on a single year (the temporally closest community matrix) seemed to perform disproportionately well, which would make sense if the community changes in a Markov-like process. We further examined the limits of predictability in terms of both hindcasting and forecasting pathogen outbreak and emergence events by training the model on a known period of 10 years, and then either forecasting or hindcasting t years into the past or future (Figure 3). Interestingly, our accuracy - measured as area under the receiver operating characteristic - did not decline at the same rate when hindcasting and forecasting. That is, model accuracy was higher when hindcasting relative to the accuracy of forecasts of the same duration of time away from the training data (Figure 3). This perhaps indicates that as the country-pathogen network becomes asymptotically more connected and stable (21), the network accumulates information content, reducing the time sensitivity of hindcasting performance.

Examining a rolling window of *t* years (*t* = 4 years) over the last two decades, we failed to detect evidence that the enhanced reporting and surveillance in more recent years influenced our model’s predictive ability (Figure 4). This also suggests that even though there were annual variations in the sample size of both pathogens and countries, there was still consistency in the structure of the country-pathogen interaction matrix over time. We explore the sensitivity of this finding to the size of the rolling window in the Supplemental Materials.

Differences in PNP model accuracy among pathogen types existed when examining the effect of the amount of data used for model training (Figure 2), with viruses having lower accuracy relative to bacteria, fungi, or other parasites. The simplest explanation for this is that accuracy is sensitive to the number of events. However, the average number of viral occurrences over time 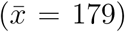 was only slightly less than the average number of bacterial 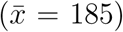 occurrences, and far greater than the average number of fungal 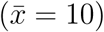 or macroparasite 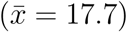 or protozoan parasite 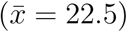 occurrence events. The average number of pathogen occurrences over time is qualitatively proportional to the number of unique viruses (*n* = 83), bacteria (*n* = 81), fungi (*n* = 14), macroparasites (*n* = 38), and protozoans (*n* = 15) we examined. Interestingly, differences among pathogen types were not found when examining the ability of the modeling approach to hind-cast/forecast (Figure 3) or when examining predictive accuracy along a rolling window (Figure 4).

For our 2016 explanatory PNP model, differentiating pathogens based on zoonotic and vector-borne transmission modes suggested that both classes of pathogens were more difficult to forecast (Figure 2). While it is possible that class imbalances between groups might drive this pattern (i.e., more event occcurences may increase model predictive accuracy), this seems unlikely: the majority of pathogens (144 of 228) were zoonotic, and many (59 of 233) were vector-borne. A more compelling explanation is that this year was an anomalous result; transmission mode did not influence accuracy when hindcasting/forecasting (Figure S5) or when models were trained along a rolling window (Figure 4), though there was notable year-to-year variation in the latter.

## Discussion

Community ecology and biogeography have a history as deeply linked fields, and both play an increasingly significant role in emerging infectious disease research. (35; 19; 36) However, research connecting the two for global pathogen diversity is fairly limited so far. Our goal was to examine whether the intrinsic structure of pathogen biogeography, approached as a bipartite network, was predictable enough to enable forecasting of different outbreak types—even in the absence of any other mechanistic predictors, like transmission mode, phylogenetic data, or environmental covariates.

Despite obvious stochasticity and data limitations, the modeling approach performed well with as little as 7 to 10 years of training data, and when predicting country-pathogen network structure across large time windows. The model was able to capture pathogen outbreak and re-emergence potential well, suggesting that, at least at administrative levels, pathogen outbreak and re-emergence events are both recurrent and predictable (and that community assembly patterns are structured and predictive of outbreak potential). However, our model generally failed to forecast pathogen emergence events. This is maybe unsurprising, as predicting when and where the next major public health threat will emerge is an incredibly difficult task which remains unsolved despite having received decades of attention (7; 1; 6). However, the failure of community information to help anticipate local emergences is still disappointing, especially given the proposal that biogeographic “co-zones” could be useful strategic tools for pandemic forecasting. (16)

We found some indications of differences in the predictability of pathogen events as a function of pathogen type and transmission modes. In the 2016 model breakdown, bacteria were the most predictable while viruses were disproportionately unpredictable, as were zoonotic and vector-borne pathogens. Given how clearly unpredictable emergence events were, this might make intuitive sense: zoonotic pathogens make up the majority of emerging diseases (1), and singlestranded RNA viruses (many vector-borne) have been responsible for many of the biggest recent emergence events (8). However, this pattern did not appear to hold up across all or even most years, and the factors that reduce model performance on a year-by-year basis are mostly unclear at the community level.

One contributor to interannual variation is large-scale events such as pandemics, which appeared to strongly influence prediction of the entire country-pathogen network. While pandemic spread may be predictable using detailed information on climate, human movement, and local environmental suitability (6; 37; 38), our approach lacks these mechanistic predictors and is sensitive to these black swan events. This can be seen in reduced model performance during the 2009 flu pandemic, including for pathogens with no relationship to flu, although viruses and vector-borne pathogens are more severely affected (see Supplemental Materials). So while the model benefits from pathogen community data, rare and widespread events can strongly reduce model accuracy. Future work to differentially weight these stochastic events would probably improve model performance.

While this approach enhances estimation of outbreak and emergence potential for rare pathogens or poorly sampled countries, it is also worth nothing that our approach is *not* a valid standalone forecasting tool. This is in large part due to how time is used in the model: though year is a covariate, the model itself is not temporally explicit, meaning that the model can predict a certain link following on previous years, but it would be erroneous to interpret that as a forecast for a given point in time. However, the tool can be used to investigate pathogen outbreak and emergence potential under different pathogen range expansion scenarios. That is, researchers could construct artificial data which differs from empirical data slightly, and quantify the ability of the model to predict those novel events. Since the method is based on dissimilarity of countries and pathogen distributions at its core, it is possible to examine the expected outcome as pathogen distributions become more (or less) homogeneous, or countries become more (or less) dissimilar in their pathogen communities.

Within infectious disease ecology, a disproportionate focus has emerged on the drivers and predictability of emergence events. (7) Recent work offers a compelling case that community ecology might bring predictive tools to bear on that problem (35), and modeling work suggests that community assembly data can be leveraged to better predict how pathogens spread (19), the host range of emerging diseases (39; 8), and the dynamics of diseases within an ecosystem (40; 41). Our results show how a simple model considering the entire pathogen community captures important global geographic variation in outbreak potential, but as a standalone tool, still struggles to predict when a pathogen will first arrive in a new region. Though this casts doubt on biogeographic tools like “co-zones” as standalone tools for surveillance or outbreak response, our study is a compelling indicator that community data could be very easily leveraged alongside other socioecological predictors to forecast disease emergence as an ecosystem process rather than a single-species one. With a Nipah virus outbreak in India and an Ebola virus outbreak in the Democratic Republic of the Congo alone both concurrent to the completion of this manuscript, the priority of prediction in emerging disease research only continues to grow.

## Acknowledgements

We thank GIDEON (https://www.gideononline.com/) for their collection and curation of the data.

Author contributions
TD, CJC, and TP conceived of the idea for the study. TD and TP designed the model. All authors contributed to the writing of the manuscript.

